# Automated compartment-based quantification reveals distinct patterns of cutaneous innervation with aging and microbial colonization

**DOI:** 10.64898/2026.10.12.758665

**Authors:** Henry Le Chang, Max Lee, Lauren Schneider, Élora Midavaine, Beatriz C. Moraes, Crystal A. Chu, Sakeen W. Kashem

## Abstract

Cutaneous innervation is commonly quantified using measures such as intraepidermal nerve fiber density, but these approaches capture only selected anatomical regions or features of nerve structure. We developed an automated image-analysis pipeline to quantify nerve area and nerve-skeleton length across defined epidermal and dermal compartments in whole, stitched mouse plantar skin sections. Automated epidermis–dermis boundary detection showed agreement with manual tracing within the range of inter-observer variation and produced similar downstream measurements. We then applied the pipeline to skin from mice across age and distinct microbial environmental exposures. Conventionalized mice exposed to microbes had greater nerve area and length than germ-free mice in both the epidermis and dermis. In the aging cohort, aged mice had greater dermal nerve area and length than young mice, while epidermal measurements did not differ significantly. We also analyzed the epidermis and dermis together to assess whole-skin innervation. Whole-skin nerve area and length were higher in conventionalized mice than in germ-free mice but did not differ significantly between young and aged mice. These findings show that analyzing the epidermis and dermis separately helps identify where nerve measurements differ, including differences that may not be detected when both skin layers are analyzed together. Compartment-resolved quantification therefore provides a complementary approach for studying changes in cutaneous innervation across biological conditions and behavioral outcomes.

## Introduction

Cutaneous nerves reside in the epidermis and dermis and are essential for somatosensation. Changes in skin innervation occur in inflammatory skin conditions, somatosensory disorders, and across development,^1, 2^ making its quantification useful in both basic and clinical research. Intraepidermal nerve fiber density (IENFD) is a widely used measure of small-fiber innervation in skin biopsies and is defined as the number of fibers crossing the epidermis–dermis boundary per millimeter of epidermal length.^3–5^ However, IENFD does not measure dermal nerves or capture changes in epidermal nerve morphology, such as nerve length or size. Complementary measurements such as fluorescent nerve area and nerve length have therefore been used in small-fiber neuropathy, aging, and nerve injury.^6–9^ Dermal nerve length has also been measured directly in human skin and can provide information not captured by IENFD.^10^

Several semi-automated methods have been developed for IENFD quantification.^11–14^ Other approaches quantify fluorescent nerve area after epidermal and dermal regions are delineated,^6^ or automate epidermal and epidermis–dermis segmentation for other skin analyses.^15, 16^ However, there is currently no end-to-end pipeline that combines automated anatomical segmentation with nerve-signal quantification across large skin samples covering both epidermal and dermal components. Such an approach could allow different features of cutaneous innervation to be compared across anatomically defined regions within the same tissue.

Microbial colonization may influence peripheral sensory systems. Germ-free (GF) mice are maintained without detectable laboratory microorganisms and can be conventionalized (ConvD) through maternal exposure to resident microbiota during pregnancy.^17, 18^ GF mice show altered inflammatory nociception and reduced maturation of the enteric nervous system, both of which can be modified by colonization.^19, 20^ Abdullah et al. found reduced nociceptor function in GF mice with effects that depended on colonization timing, but no difference in nerve fiber length in either the epidermis or dermis.^21^ Under other experimental conditions, microbiota-driven immune responses have been associated with sensory-neuron regeneration and skin hyperinnervation.^22, 23^ It therefore remains unclear whether compartment-resolved measurements can identify and anatomically localize differences among mice with and without microbiome exposure that may not be captured by previously used fiber-length measurements.

Aging provides one setting in which the choice of measurement may be important. Several studies report reduced IENFD in aged human and mouse skin,^24–27^ whereas others report minimal age-related change or nonlinear changes across the lifespan.^8, 28^ Epidermal fluorescent nerve area has also been reported to decrease, remain unchanged, or increase with age depending on anatomical site.^7^ In addition, Metzner et al. reported increased cytoskeletal staining intensity in sensory axons and increased axon diameter in aged males but not females.^29^ Behavioral responses are similarly variable: mechanical sensitivity generally decreases in aged mice, while responses to nociceptive stimuli depend on modality, sex, and experimental conditions.^30^ Together, these findings suggest that different measurements and anatomical compartments may capture different age-associated changes in cutaneous innervation.

We developed an automated image-analysis pipeline that segments skin into anatomical compartments and quantifies *β*III-tubulin (TUBB3)-positive nerve area and skeleton length. We validated the automated epidermis–dermis boundary against manual traces and applied the pipeline to plantar skin from GF and ConvD mice and from young and aged mice. In the microbiota cohort, ConvD mice showed higher epidermal, dermal, and whole-skin measurements than GF mice, with significant differences in the deep dermis but not the superficial dermis. In the aging cohort, the clearest differences were dermal, including significant dermal differences despite nonsignificant whole-skin measurements. These findings illustrate how compartment-resolved analysis can identify dermal differences despite nonsignificant whole-skin comparisons and map differences spanning epidermal and dermal compartments.

## Results

### Automated compartment-based analysis quantifies axonal cutaneous innervation

We acquired stitched images of longitudinal mouse plantar skin sections stained for DAPI and *β*III-tubulin (TUBB3) (Figure 1A). DAPI served as an anatomical landmark, and an ilastik pixel-classification model was used to generate lower-epidermal and whole-skin masks from the DAPI channel (Figure 1B).^31^ These masks were processed to define four mutually exclusive compartments: upper epidermis, lower epidermis, superficial dermis, and deep dermis (Figure 1C). The superficial (subepidermal) dermis was defined as the region within a 20-*µ*m geodesic distance below the derived epidermis–dermis boundary, and the remaining dermis was classified as deep dermis.

The anatomical compartments were applied to the TUBB3 channel after background subtraction and thresholding (Figure 1D). The pipeline measured TUBB3-positive area and skeletonized the segmented signal to measure skeleton length (Figure 1E). Upper and lower epidermis were combined for whole-epidermis measurements, while superficial and deep dermis were combined for whole-dermis measurements. Epidermal TUBB3-positive area and skeleton length were normalized to epidermis–dermis boundary length; dermal and whole-skin measurements were normalized to tissue area, giving TUBB3-positive area fraction and skeleton density. We also measured TUBB3-positive area per skeleton length in the superficial dermis as a measure related to the apparent width of segmented TUBB3-positive profiles.^29^

**Figure 1.**
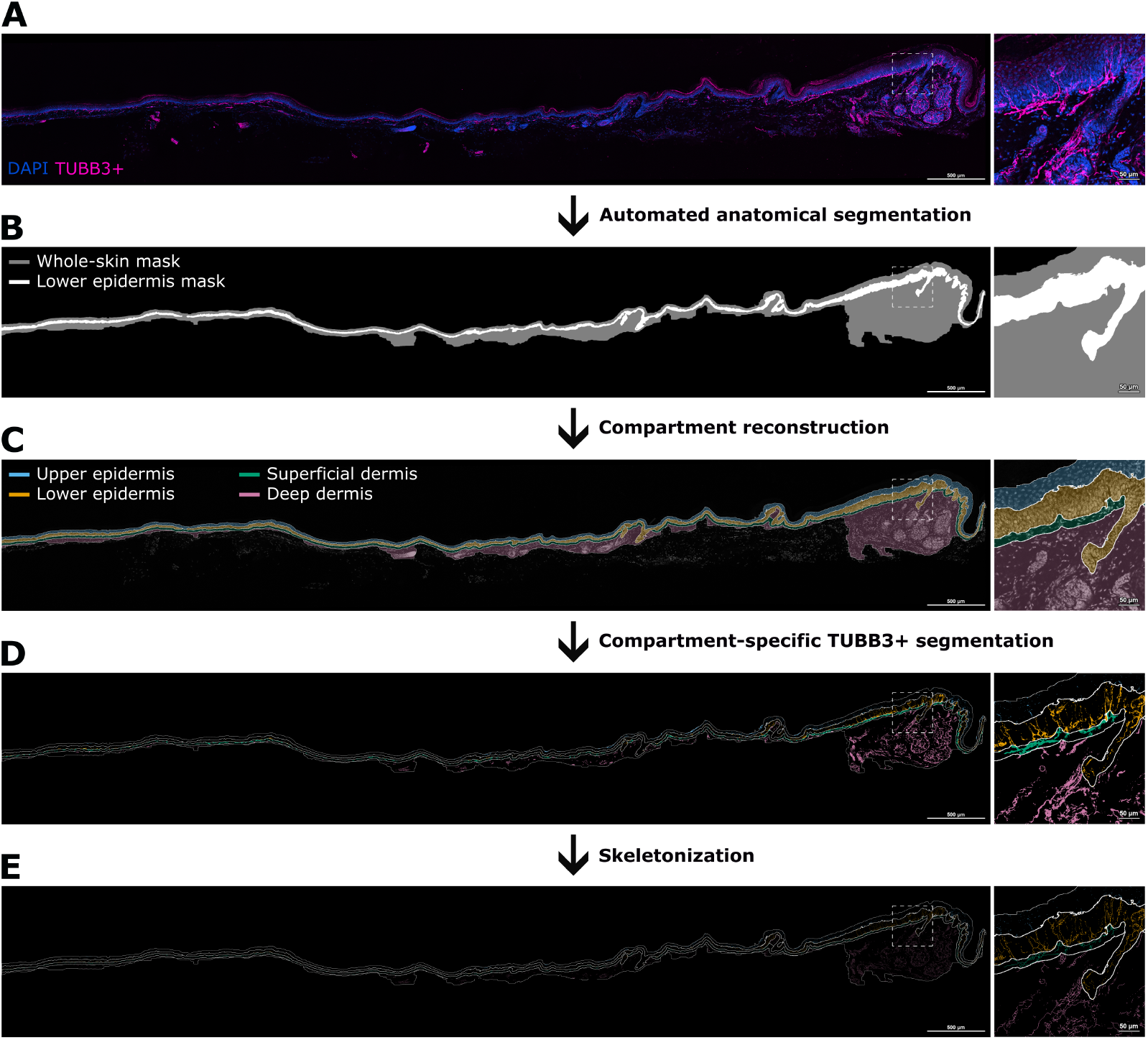
Automated compartment-based analysis quantifies TUBB3-positive cutaneous innervation. (A) Representative stitched maximum-intensity projection of a longitudinal mouse plantar skin section stained for DAPI (blue) and TUBB3 (magenta), with a magnified region shown at right. (B) Automated whole-skin mask (gray) and lower-epidermal mask (white) generated from the DAPI channel. (C) Reconstruction of four anatomical compartments: upper epidermis, lower epidermis, superficial dermis, and deep dermis. The superficial dermis was defined as dermal tissue within 20 *µ*m geodesic distance below the derived epidermis–dermis boundary. Upper and lower epidermis were combined for whole-epidermis measurements, superficial and deep dermis were combined for whole-dermis measurements, and all four compartments comprised the whole-skin region. (D) Thresholded TUBB3-positive signal assigned to the corresponding anatomical compartments. (E) Skeletonized TUBB3-positive signal used for length-based measurements. Epidermal measurements were normalized to epidermis–dermis boundary length; dermal and whole-skin measurements were normalized to tissue area. Scale bars, 500 *µ*m in whole-section images and 50 *µ*m in magnified regions.

### Automated epidermal boundary detection agrees with manual tracing

To validate automated boundary detection, we compared the automated dermal-epidermal junction boundary with manually traced boundaries from two observers using eight held-out sections from four mice (Figure 2A). The automated segmentation used DAPI; CD49f fluorescence was available only as an additional anatomical guide during manual tracing. Agreement was assessed using average symmetric surface distance (ASSD) and the Surface Dice similarity coefficient with a 10-*µ*m tolerance.^32, 33^

Mean ASSD was 5.62 *µ*m between the automated boundary and observer 1, 3.63 *µ*m between the automated boundary and observer 2, and 4.32 *µ*m between observers 1 and 2 (Figure 2B). Mean Surface Dice was 90.0%, 95.5%, and 93.0%, respectively (Figure 2C). Thus, automated–manual boundary agreement was within the range of inter-observer agreement.

**Figure 2.**
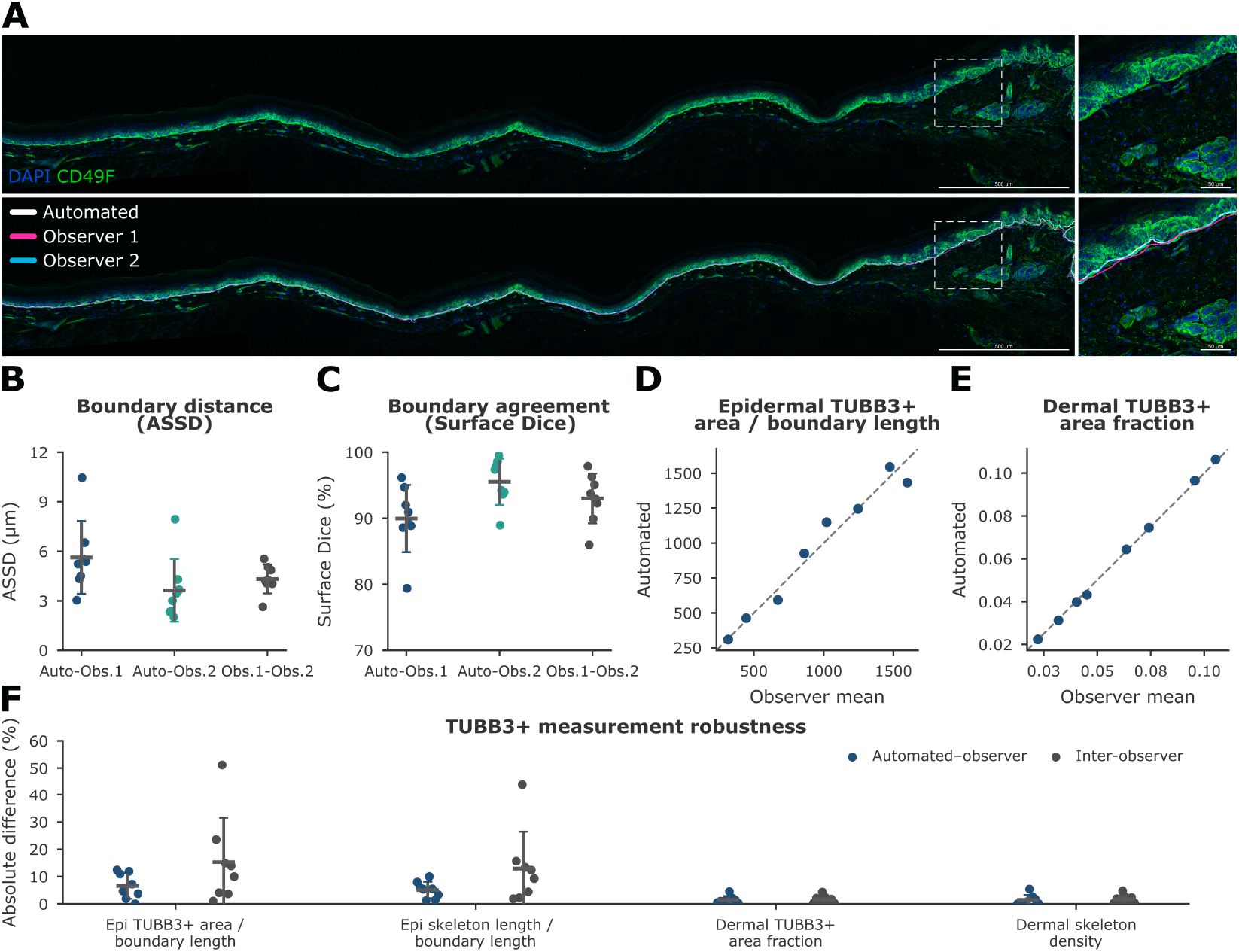
Automated epidermis–dermis boundary detection agrees with manual tracing. (A) Representative longitudinal plantar skin section stained for DAPI (blue) and CD49f (green), with the automated epidermis–dermis boundary and boundaries traced by two independent observers overlaid below. Magnified regions are shown at right. (B) Average symmetric surface distance (ASSD) between the automated boundary and observer 1, the automated boundary and observer 2, and the two observer-defined boundaries. (C) Surface Dice similarity coefficient using a 10-*µ*m tolerance for the same comparisons. (D–E) Comparison of TUBB3-positive measurements generated using the automated boundary and the mean of the observer-defined boundaries for epidermal TUBB3-positive area per boundary length and dermal TUBB3-positive area fraction. Dashed lines indicate identity. (F) Absolute percentage differences between automated and observer-defined measurements compared with inter-observer differences for epidermal TUBB3-positive area per boundary length, epidermal skeleton length per boundary length, dermal TUBB3-positive area fraction, and dermal skeleton density. Validation included eight sections from four mice, with two sections from each mouse. Points represent section-level comparisons and bars show mean ± SEM. Scale bars, 500 *µ*m in whole-section images and 50 *µ*m in magnified regions.

We also tested whether boundary placement affected downstream TUBB3-positive measurements. Mean absolute differences between automated and observer-derived measurements were 6.61% for epidermal TUBB3-positive area per boundary length, 5.11% for epidermal nerve skeleton length per boundary length, 1.48% for dermal TUBB3-positive area fraction, and 1.47% for dermal skeleton density. The corresponding inter-observer differences were 15.30%, 12.90%, 1.66%, and 1.72%, respectively (Figure 2D–F). Overall, automated boundary placement was within the range of manual inter-observer variation and produced similar downstream TUBB3-positive measurements.

### Commensal microbes coordinate cutaneous innervation

To investigate whether microbial colonization status was associated with skin innervation, we analyzed plantar skin from five GF and five ConvD female mice (ten mice total; Figure 3A). Two sections from each mouse were analyzed, giving ten sections per group. Section-level measurements were averaged within each mouse, yielding five biological replicates per group. Groups were compared using two-sided unpaired Welch’s t-tests, with P values unadjusted across the ten histological outcomes.

**Figure 3.**
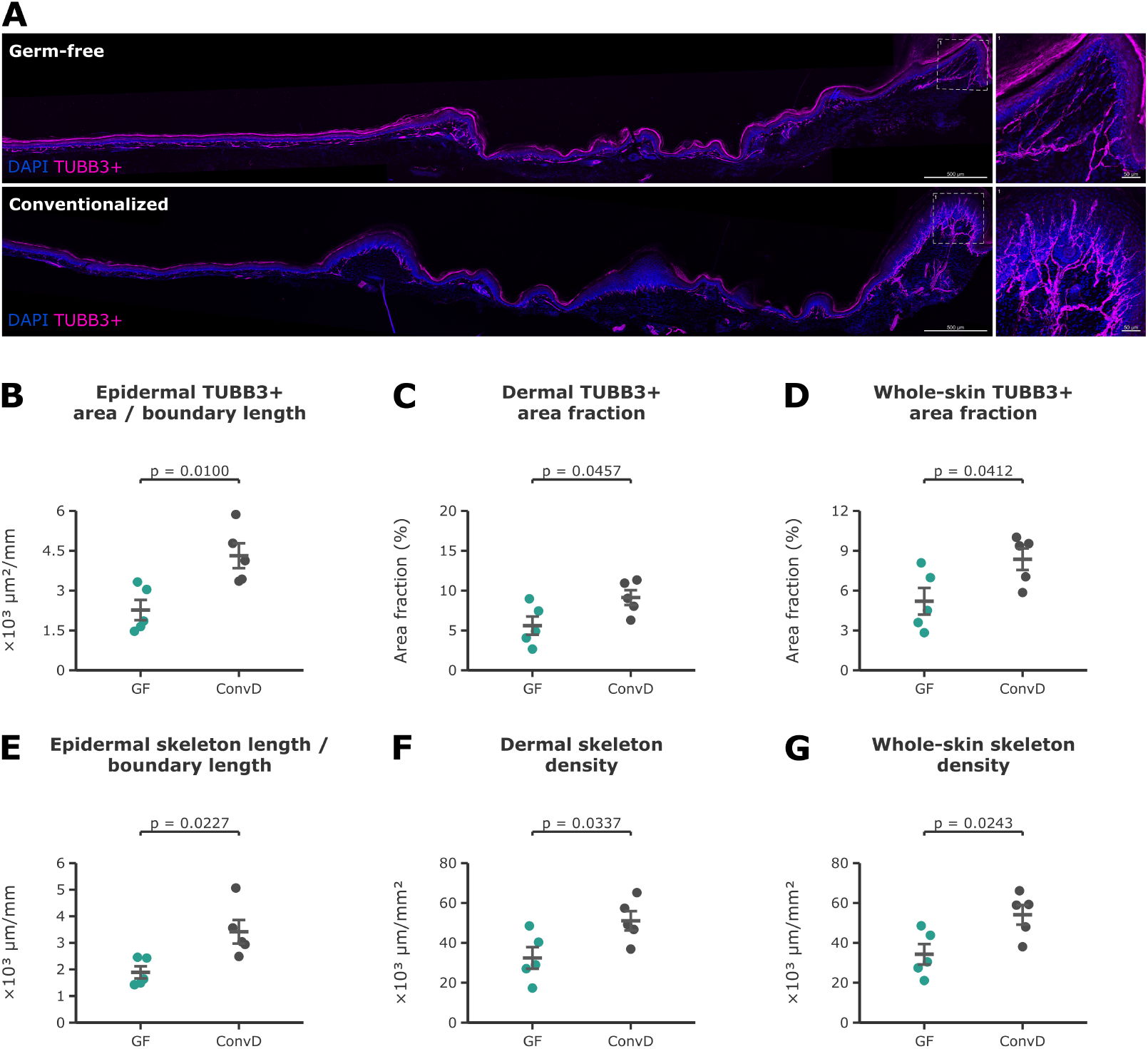
Conventionalized mice show higher epidermal, dermal, and whole-skin TUBB3-positive measurements. (A) Representative stitched longitudinal plantar skin sections from germ-free (GF) and conventionalized (ConvD) female mice stained for DAPI (blue) and TUBB3 (magenta), with magnified regions shown at right. (B) Epidermal TUBB3-positive area per epidermis–dermis boundary length. (C) Dermal TUBB3-positive area fraction. (D) Whole-skin TUBB3-positive area fraction. (E) Epidermal skeleton length per epidermis–dermis boundary length. (F) Dermal skeleton density. (G) Whole-skin skeleton density. GF and ConvD groups each included n = 5 female mice, with two sections averaged per mouse. Comparisons were performed using two-sided unpaired Welch’s t-tests. Points represent individual mice and bars show mean ± SEM. P values are shown and are unadjusted across the ten GF-versus-ConvD histological outcomes. Scale bars, 500 *µ*m in whole-section images and 50 *µ*m in magnified regions.

ConvD mice had significantly higher epidermal neurite-positive area per boundary length than GF mice (4.31 ± 0.47 vs. 2.27 ± 0.38 × 10^3^ *µ*m^2^/mm, P = 0.0100; Figure 3B). Epidermal skeleton length per boundary length was also higher in ConvD mice (3.42 ± 0.45 vs. 1.89 ± 0.23 × 10^3^ *µ*m/mm, P = 0.0227; Figure 3E).

Dermal neurite-positive area fraction was higher in ConvD mice than GF mice (9.12 ± 0.93% vs. 5.61 ± 1.15%, P = 0.0457; Figure 3C), as was dermal skeleton density (51.04 ± 4.81 vs. 32.42 ± 5.43 × 10^3^ *µ*m/mm^2^, P = 0.0337; Figure 3F). Deep-dermal neurite-positive area fraction (8.12 ± 0.94% vs. 4.81 ± 1.00%, P = 0.0427; Figure S1G) and skeleton density (44.96 ± 5.10 vs. 27.79 ± 4.72 10^3^ *µ*m/mm^2^, P = 0.0388; Figure S1H) were also higher in ConvD mice. In contrast, superficial neurite-positive area fraction (13.74 ± 1.52% vs. 9.69 ± 1.97%, P = 0.1435; Figure S1E) and skeleton density (77.98 ± 5.97 vs. 55.44 ± 9.65 10^3^ *µ*m/mm^2^, P = 0.0894; Figure S1F) did not differ significantly between groups.

To assess overall skin innervation, we combined the epidermal and dermal regions and normalized nerve-positive area and skeleton length to the total tissue area. ConvD mice also had higher whole-skin nerve-positive area fraction than GF mice (8.36 ± 0.81% vs. 5.20 ± 1.00%, P = 0.0412; Figure 3D) and higher whole-skin skeleton density (54.05 ± 4.95 vs. 34.26 ± 5.15 × 10^3^ *µ*m/mm^2^, P = 0.0243; Figure 3G). Thus, the GF–ConvD differences were detected in both epidermal and dermal measurements, with significant differences in the deep dermis but not the superficial dermis.

### Aging disrupts sensory nervous system architecture and function

To investigate age-associated differences in skin innervation, we analyzed plantar skin from seven young male mice and six aged male mice (Figure 4A). Three sections from each mouse were analyzed and averaged so that each mouse contributed one biological replicate. Young-versus-aged comparisons were performed using two-sided unpaired Welch’s t-tests.

**Figure 4.**
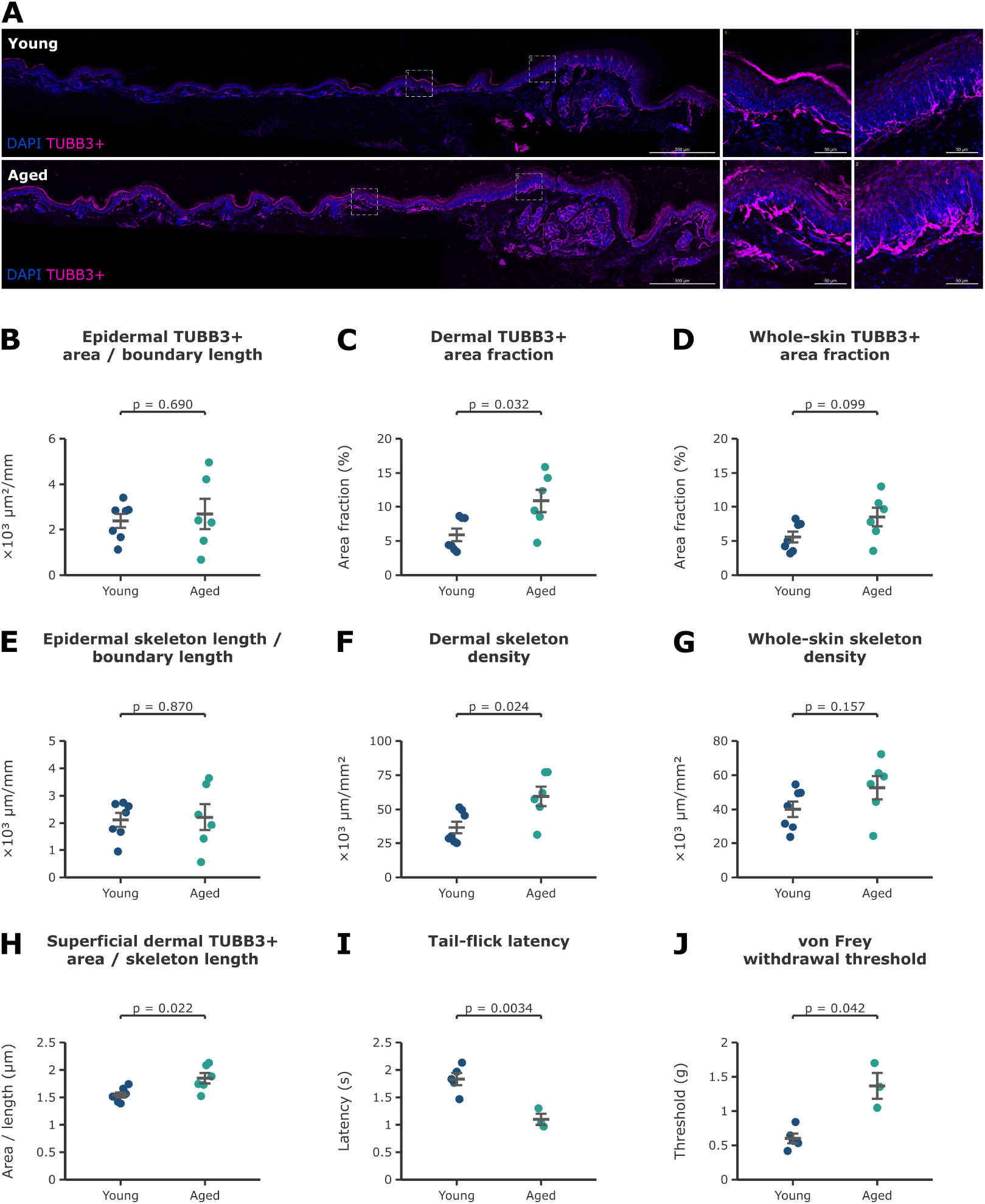
Age-associated differences in TUBB3-positive measurements are most apparent in the dermis. (A) Representative stitched longitudinal plantar skin sections from young and aged male mice stained for DAPI (blue) and TUBB3 (magenta), with two magnified regions shown at right for each group. (B) Epidermal TUBB3-positive area per epidermis–dermis boundary length. (C) Dermal TUBB3-positive area fraction. (D) Whole-skin TUBB3-positive area fraction. (E) Epidermal skeleton length per epidermis–dermis boundary length. (F) Dermal skeleton density. (G) Whole-skin skeleton density. (H) TUBB3-positive area per skeleton length in the superficial dermis, a measurement related to the apparent width of segmented TUBB3-positive profiles. Histological measurements were obtained from seven young mice and six aged mice, with three sections averaged per mouse. (I) Tail-flick latency and (J) von Frey withdrawal threshold in a subset of five young and three aged mice. Young and aged groups were compared using two-sided unpaired Welch’s t-tests. Points represent individual mice and bars show mean ± SEM. Exact P values are shown. Scale bars, 500 *µ*m in whole-section images and 50 *µ*m in magnified regions.

Aged mice had significantly higher dermal nerve-positive area fraction than young mice (10.87 ± 1.67% vs. 5.89 ± 0.91%, P = 0.032; Figure 4C) and higher dermal skeleton density (59.53 ± 7.07 vs. 36.64 ± 4.43 × 10^3^ *µ*m/mm^2^, P = 0.024; Figure 4F). In contrast, epidermal nerve-positive area per boundary length (2.68 ± 0.66 vs. 2.38 0.31 × 10^3^ *µ*m^2^/mm, P = 0.690; Figure 4B) and epidermal skeleton length per boundary length (2.21 ± 0.48 vs. 2.12 ± 0.26 × 10^3^ *µ*m/mm, P = 0.870; Figure 4E) did not differ significantly.

Nerve-positive area fraction in the superficial dermis was also higher in aged mice (16.54 ± 2.38% vs. 8.36 ± 1.50%, P = 0.018; Figure S1A), while deep-dermal area fraction did not differ significantly (P = 0.052; Figure S1C). Skeleton density was higher in aged mice in both the superficial dermis (87.17 ± 8.72 vs. 52.65 ± 8.16 × 10^3^ *µ*m/mm^2^, P = 0.015; Figure S1B) and deep dermis (51.18 ± 6.69 vs. 32.17 ± 3.76 10^3^ *µ*m/mm^2^, P = 0.038; Figure S1D). Aged mice also had higher nerve-positive area per skeleton length in the superficial dermis (1.85 ± 0.09 vs. 1.54 ± 0.05 *µ*m, P = 0.022; Figure 4H), consistent with greater apparent width of segmented neurite-positive profiles, although this measurement does not directly measure individual axon diameter.

Although aged mice had higher dermal nerve-positive area fraction and skeleton density, neither measure differed significantly when the epidermis and dermis were analyzed together (P = 0.0992 and 0.1570, respectively; Figure 4D,G). Thus, the clearest age-associated histological differences were dermal.

We also assessed sensory behavior in a subset of young and aged mice. Aged mice had a shorter tail-flick latency than young mice (1.10 ± 0.10 vs. 1.83 ± 0.11 s, *P* = 0.0034; Figure 4I), indicating greater responsiveness to noxious heat, and a higher von Frey withdrawal threshold (1.37 ± 0.19 vs. 0.60 ± 0.07 g, *P* = 0.0416; Figure 4J), indicating reduced mechanical sensitivity.

## Discussion

ConvD mice had greater nerve-positive area and skeleton length than GF mice in the epidermis, dermis, and whole skin. Within the dermis, deep-dermal measurements differed significantly, whereas superficial measurements did not. In the aging cohort, aged mice had higher dermal neurite-positive area fraction and skeleton density than young mice, with no significant differences in epidermal or whole-skin measurements. Together, these findings illustrate two uses of compartment-resolved analysis: resolving the anatomical distribution of higher whole-skin measurements in the microbiota cohort and identifying dermal differences despite nonsignificant whole-skin measurements in the aging cohort.

The GF–ConvD differences in nerve-positive area and skeleton length spanned both the epidermis and dermis. Previous studies have shown altered neural function or maturation in GF animals compared with colonized animals.^19, 34^ Abdullah et al. found no difference in skin nerve fiber length among microbial colonization groups despite differences in nociceptor sensitivity.^21^ Their colonization paradigm is not directly equivalent to ours, and they quantified fiber length in sampled plantar-skin regions rather than neurite-positive area and skeleton length across entire stitched sections. Differences in anatomical sampling, quantification method, colonization timing, or microbiota composition across facilities may therefore contribute to the different findings.^35, 36^ Microbiota-driven immune responses have also been linked to peripheral nerve regeneration and skin hyperinnervation under other experimental conditions.^22, 23^ Future experiments using defined microbial communities could help determine which microbial exposures, if any, contribute to the epidermal and dermal differences observed in ConvD mice.

Much of the literature reports reduced IENFD with age,^24, 25, 27^ whereas our epidermal measurements did not differ significantly between groups. This may reflect differences in what each measurement captures. Epidermal fluorescent nerve area has been reported to decrease, remain unchanged, or increase with age depending on biopsy site,^7^ and Lauria et al. found relatively little age-related change in epidermal innervation in healthy individuals up to approximately 75 years of age.^28^ In rat plantar skin, both IENFD and epidermal nerve length showed a nonlinear relationship with age, increasing through middle age before declining later.^8^ Thus, area- and skeleton-based measurements may capture age-associated changes differently from IENFD.

Higher dermal nerve-positive area fraction and skeleton density in aged mice were accompanied by a higher nerve-positive area per skeleton length in the superficial dermis, a measure related to apparent profile width. Previous studies of dermal innervation with age are mixed. Kelly et al. reported reduced papillary dermal nerve fiber density with age in human glabrous hand skin despite little corresponding change in epidermal innervation.^37^ Gavini et al. found fewer sub-basal nerve plexus profiles in aged mouse plantar skin, but quantified PGP9.5-positive profiles per unit length rather than nerve area or length.^27^ In contrast, Metzner et al. reported increased TUBB3 immunostaining intensity and increased sensory-axon diameter specifically in aged human males.^29^ Their measurements are not equivalent to ours, but the higher TUBB3 signal is directionally consistent with the higher dermal nerve-positive area fraction observed in aged mice in our cohort. Their area-to-length approach for estimating axon diameter is also conceptually related to our nerve-positive area per skeleton length measurement, which was higher in aged male mice. Together, these studies suggest that apparent age-associated changes depend on anatomical compartment, sex, and how innervation is quantified.

In the aging cohort, aged mice showed reduced mechanical sensitivity, consistent with previous reports, while thermal responses with aging vary across assays and experimental conditions.^8, 30, 38^ Overall, we hypoth-esize that these age-associated changes may potentially reflect axonal edema and peripheral neurodegeneration that link neuronal anatomy to altered sensory behavioral output. Studies spanning both sexes and multiple ages will be needed to determine how nervous system innervation and behavior correlate across the lifespan.

Several limitations should be considered. The analysis uses two-dimensional maximum-intensity projections of thick skin sections, which simplify three-dimensional nerve organization and can merge overlapping nerve-positive profiles; skeleton length therefore does not represent total three-dimensional axonal length. The aging cohort contained only male mice, whereas the microbiota cohort contained only female mice, so the cohorts should not be compared directly and the findings may not generalize across sexes. Microbiota composition can also vary among facilities and may contribute to differences between GF and ConvD groups across different studies and sites. In addition, thresholded TUBB3-positive area and skeleton measurements quantify labeled signal rather than individual fibers and can be influenced by staining intensity, profile caliber, or signal continuity. The GF–ConvD comparison included five mice per group, and P values were unadjusted across ten histological outcomes; confirmation in independent cohorts is therefore needed. Finally, the cross-sectional design does not establish how these differences develop over time or whether they cause the measured sensory responses.

Overall, compartment-resolved analysis distinguished the anatomical distribution of nerve-positive changes across cohorts. In ConvD mice, higher whole-skin measurements were accompanied by higher epidermal and dermal measurements, with significant differences in the deep dermis but not the superficial dermis. In the aging cohort, the clearest age-associated differences were dermal despite nonsignificant whole-skin measurements. These results support compartment-based quantification as a complementary approach for revealing region-specific changes and anatomically localizing broader changes in cutaneous innervation and sensory function.

## Methods

### Experimental animals and cohorts

The aging cohort consisted of seven young adult (7–8-week-old) and six aged (19–24-month-old) male C57BL/6J mice. Young adult mice were purchased from The Jackson Laboratory. Of the six aged mice, three were obtained from The Jackson Laboratory and three were provided by the laboratory of Dr. Saul Villeda at UCSF. All young and aged mice were maintained at the University of California, San Francisco (UCSF) Laboratory Animal Resource Center, Mission Bay campus. Germ-free mice were provided by the UCSF gnotobiotic facility. Pregnant germ-free mice were exposed to germ-free stool and bedding (GF) or stool and bedding from mice maintained under specific pathogen-free conditions (ConvD). Their offspring were used for experiments. The microbiota histology analysis included five GF and five ConvD female mice, with plantar skin collected at 8 weeks of age. All animal procedures were approved by the UCSF Institutional Animal Care and Use Committee.

### Plantar skin preparation, staining, and imaging

Plantar skin was fixed, cryoprotected, sectioned longitudinally at 50 *µ*m, and stained for TUBB3, CD49f, and DAPI. Whole sections were imaged as tiled confocal z-stacks and stitched for analysis. TUBB3 images underwent rolling-ball background subtraction before maximum-intensity projection. Detailed tissue preparation, staining, imaging, and preprocessing procedures are provided in Supplementary Methods.

### Automated anatomical segmentation and TUBB3 quantification

DAPI images were analyzed using ilastik v1.4.2^31^ to segment the lower-epidermal DAPI band and whole skin. Fixed postprocessing generated four mutually exclusive compartments: upper epidermis, lower epidermis, superficial dermis, and deep dermis. The superficial dermis was defined as dermal tissue within 20 *µ*m geodesic distance of the derived epidermis–dermis boundary; the remaining dermis was classified as deep dermis. Upper and lower epidermis were combined for whole-epidermis measurements, superficial and deep dermis were combined for whole-dermis measurements, and all four compartments comprised the whole-skin region.

For TUBB3 quantification, a fixed intensity threshold of > 1500 was applied to every preprocessed image.

Connected TUBB3-positive objects with a skeleton length of at least 20 *µ*m were excluded when any portion was within 5 *µ*m of the outer skin surface. The remaining signal was restricted to the whole-skin mask. TUBB3-positive area and skeleton length were measured within each compartment. Epidermal measurements were normalized to epidermis–dermis boundary length, whereas dermal and whole-skin measurements were normalized to the corresponding tissue area. TUBB3-positive area per skeleton length was additionally calculated in the superficial dermis as a measure related to apparent profile width. The same preprocessing, threshold, and analysis parameters were used across samples. OpenAI Codex was used to assist with code development and debugging. All AI-generated code was reviewed, tested, and modified as needed before use.

### Validation and biological replicates

The ilastik models and downstream analysis parameters were finalized before validation. Automated epidermis–dermis boundaries were evaluated against traces from two independent observers in eight held-out full-length sections using ASSD and Surface Dice with a 10-*µ*m tolerance.^32, 33^ CD49f was not used by the automated segmentation and was available only as an anatomical guide for manual tracing. Downstream epidermal and dermal TUBB3-positive measurements were also compared between automated and observer-defined boundaries. Additional model-development and validation details are provided in Supplementary Methods.

For aging analyses, three sections per mouse were quantified. For microbiota analyses, two sections per mouse were quantified. Sections were separated by approximately 300–400 *µ*m and averaged within each mouse so that each mouse contributed one biological replicate.

### Behavioral testing

Mechanical and heat sensitivity were assessed in five young and three aged mice using von Frey filaments and the tail-flick assay, respectively. Behavioral testing procedures were adapted from Midavaine et al.^39^ Detailed procedures are provided in Supplementary Methods.

### Statistical analysis

All histological statistical analyses used mouse-level averages, with each mouse treated as one biological replicate. Young-versus-aged comparisons and GF-versus-ConvD histological comparisons used two-sided unpaired Welch’s t-tests. For GF-versus-ConvD comparisons, five mice per group were analyzed, and P values were calculated using Welch–Satterthwaite degrees of freedom without adjustment for multiple comparisons across the ten histological outcomes. Young-versus-aged behavioral comparisons also used two-sided unpaired Welch’s t-tests. Histological analyses for both cohorts were performed in Python 3.13.5 using SciPy 1.17.0. Data are presented as mean ± SEM, and P < 0.05 was considered significant.

## Data availability

Data supporting the findings of this study are included in the article and Supplementary Information. Raw imaging data, processed measurements, and additional analysis outputs are available from the corresponding authors upon reasonable request.

## Code availability

The custom Python code and ilastik models used for anatomical segmentation, TUBB3 quantification, validation, and statistical analysis are available at GitHub: https://github.com/henrylechang/Skin-Analysis. The repository will be updated as the analysis pipeline is further developed.

## Acknowledgments

We thank Saul Villeda and members of his laboratory for providing aged mice and the UCSF gnotobiotic core for germ-free and conventionalized mice used in this study. We thank Allan I. Basbaum for providing resources.

This study received support from National Institutes of Health (NIH) grant 1K99NS142248-01 (É.M.); Banting Canadian Institute of Health Research (CIHR Award No. BPF198457) (É.M.); the Fonds de Recherche en Santé-Québec (É.M.); the Sandler Foundation PBBR (É.M. and S.W.K.); the Dermatology Foundation (Career Development Award) (S.W.K.); the Benioff Microbiome Research Program (S.W.K.); a Grunfeld Scholar Award from SFVAMC (S.W.K.); National Institutes of Health (NIH) grants T32AR007175-44, R03NS148838, K08AR085766, and DP2NS155519 (S.W.K.); Arthritis National Research Foundation (S.W.K.); Burroughs Wellcome Fund Career Award for Medical Scientists (S.W.K.); ISID-LEO Foundation Young Investigator Prize (S.W.K.); and Pew Latin American Scholars (B.C.M.).

## Author contributions

H.L.C. conceived the project, performed experiments, analyzed data, and wrote the manuscript. M.L., L.S., É.M., B.C.M., and C.A.C. performed experiments. S.W.K. conceived the project, supervised the study, and contributed to writing the manuscript. All authors reviewed and approved the manuscript.

## Competing interests

The authors declare no competing interests.

## Notes

https://github.com/henrylechang/Skin-Analysis

### Competing Interest Statement

The authors have declared no competing interest.

