## Supplementary Information for "Automated compartment-based quantification reveals distinct patterns of cutaneous innervation with aging and microbial colonization"

Supplementary Methods and Figure S1

### Supplementary Methods

#### Mouse plantar skin preparation and cryosectioning

Paws were collected and immersion-fixed in 4% paraformaldehyde at 4°C for 2–3 days. Following fixation, paws were washed in phosphate-buffered saline (PBS), and plantar skin was dissected from the underlying tissue. Major ligaments and excess connective tissue were removed, and the isolated plantar skin was cryoprotected in 30% sucrose for 2–3 days.

Cryoprotected plantar skin was embedded flat in optimal cutting temperature compound and frozen on dry ice. Tissue blocks were equilibrated in the cryostat for at least 30 min before sectioning longitudinally. Sections were cut at 50 $\mu$m and distributed sequentially across seven slides so that sections collected on the same slide were separated by approximately 300–400 $\mu$m. Sections were dried at room temperature for at least 3 h before storage at -80°C.

#### Immunofluorescence staining

Frozen sections were equilibrated at room temperature for 1 h and washed twice in PBS for 10 min. Sections were blocked and permeabilized for 1 h at room temperature in PBS containing 1% Triton X-100 and 5% normal goat serum. Sections were then incubated overnight at 4°C in the same buffer containing Alexa Fluor 647-conjugated anti-$\beta$III-tubulin antibody (TUBB3; clone TUJ1; BioLegend, cat. no. 801210; 1:300), Alexa Fluor 488-conjugated anti-human/mouse CD49f antibody (clone GoH3; BioLegend, cat. no. 313608; 1:300), and DAPI (1:10,000). Following staining, sections were washed twice in PBS for 10 min and mounted using Fluoromount.

#### Confocal imaging and image preprocessing

Fluorescence images were acquired using an Olympus FLUOVIEW FV3000 laser-scanning confocal microscope with a 20× objective. Large tissue sections were imaged as tiled fields and stitched to generate whole-section images. DAPI, Alexa Fluor 488, and Alexa Fluor 647 fluorescence were acquired separately. Individual fields were acquired at 1040 × 1040 pixels as z-stacks with a 3-$\mu$m step size. Images had a pixel size of 0.621504 $\mu$m/pixel in both the aging and microbiota cohorts. Fluorescence was acquired at 12-bit depth, with intensity values ranging from approximately 0 to 4096. Imaging parameters were held constant across samples within each experimental comparison.

Multichannel z-stacks were separated into DAPI, TUBB3, and CD49f channels. Maximum-intensity projections were generated from the DAPI and CD49f stacks. For TUBB3, background subtraction was performed in Fiji (ImageJ v1.54p) using a rolling-ball algorithm with a 7.5-$\mu$m radius before maximum-intensity projection. The resulting single-channel images were saved as 16-bit grayscale TIFF files without rescaling the fluorescence intensity range and were rotated as necessary to standardize tissue orientation, with the epidermis positioned toward the top of the image. CD49f images were retained for validation of the epidermis–dermis boundary.

#### Segmentation model development and validation details

The ilastik segmentation models were developed using images from four mice: two 8-week-old mice, one 12-week-old mouse, and one 24-month-old mouse. Segmentation models and downstream image-processing parameters were finalized before evaluation of the validation dataset. Validation images were obtained from separately stained sections and were not used as training images.

Automated epidermis–dermis boundary detection was evaluated using eight full-length plantar skin sections from four mice: two young and two aged mice, with two sections from each mouse. Two independent observers manually traced the epidermis–dermis boundary using the DAPI channel, with CD49f fluorescence available as an additional anatomical guide. Agreement between automated and observer-defined boundaries was quantified using average symmetric surface distance (ASSD) and Surface Dice with a 10-$\mu$m tolerance.

To determine whether boundary placement affected downstream measurements, epidermal TUBB3-positive area per boundary length, epidermal skeleton length per boundary length, dermal TUBB3-positive area fraction, and dermal skeleton density were compared using automated and observer-defined boundaries.

#### Behavioral testing

##### von Frey test

Mice underwent a single 2-h acclimation session in the von Frey apparatus. On each test day, mice were habituated on a wire mesh for 1 h before testing. Mechanical sensitivity was measured by applying von Frey filaments (Stoelting, cat. no. 58011; 0.008, 0.02, 0.04, 0.07, 0.16, 0.4, 0.6, 1, 1.4, 2, 4, and 8 g) to the lateral plantar surface of the hind paws using the Dixon up–down method. Each filament was applied for 3 s, with several minutes between successive stimulations. The 50% paw withdrawal threshold was measured on two days, and the values were averaged.

##### Tail-flick test

Mice underwent a single acclimation session in a restrainer, during which the tail was immersed in room-temperature water. On the test day, mice were acclimated for 30 min in plexiglass cylinders before being placed in the restrainer. The distal 2 cm of the tail was submerged in a 52°C water bath, and the latency to withdraw the tail was recorded in seconds. A 15-s cutoff was imposed to prevent tissue damage. Three trials were performed, separated by several minutes, and withdrawal latencies were averaged for each mouse.

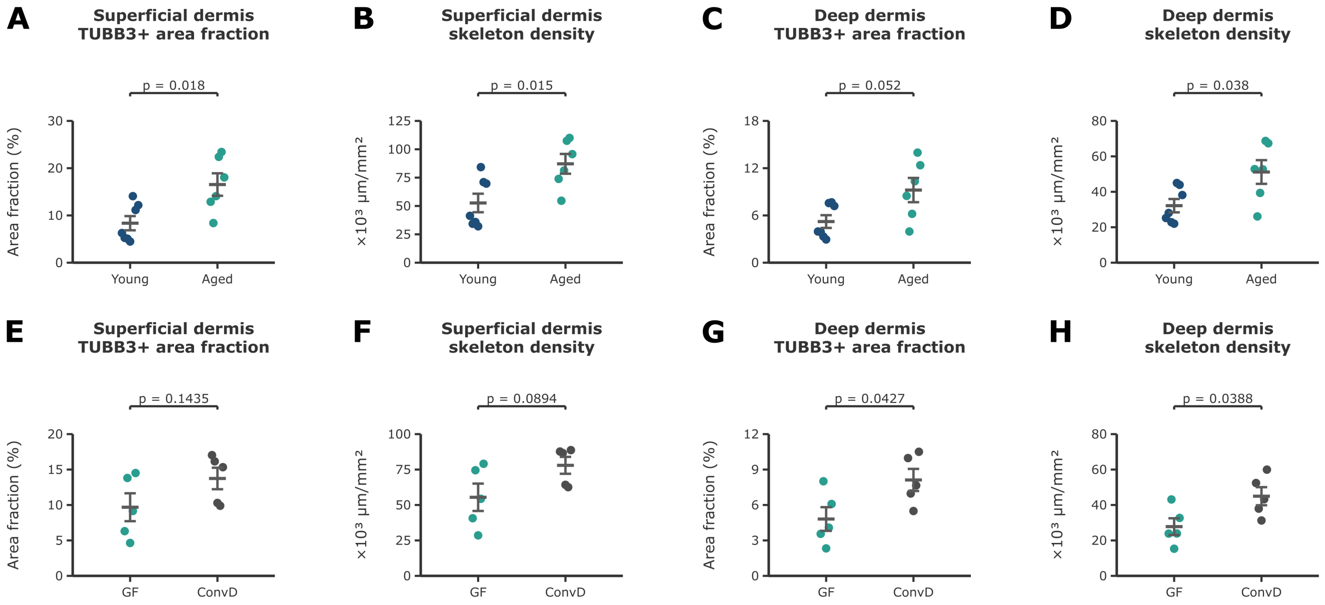


**Figure S1. Superficial- and deep-dermal TUBB3-positive measurements in aging and microbial colonization cohorts.** (A–D) Measurements from young and aged male mice. (A) Superficial dermal TUBB3-positive area fraction. (B) Superficial dermal skeleton density. (C) Deep-dermal TUBB3-positive area fraction. (D) Deep-dermal skeleton density. Seven young and six aged mice were analyzed, with three sections averaged per mouse. Young and aged groups were compared using two-sided unpaired Welch's t-tests. Superficial dermal area fraction and skeleton density and deep-dermal skeleton density were significantly higher in aged mice, while deep-dermal area fraction did not differ significantly between groups. (E–H) Measurements from germ-free (GF) and conventionalized (ConvD) female mice. (E) Superficial dermal TUBB3-positive area fraction. (F) Superficial dermal skeleton density. (G) Deep-dermal TUBB3-positive area fraction. (H) Deep-dermal skeleton density. GF and ConvD groups each included n = 5 female mice, with two sections averaged per mouse. GF and ConvD groups were compared using two-sided unpaired Welch's t-tests, with P values unadjusted across the ten GF-versus-ConvD histological outcomes. ConvD mice had significantly higher deep-dermal TUBB3-positive area fraction and skeleton density than GF mice; superficial dermal area fraction and skeleton density did not differ significantly between groups. Points represent individual mice and bars show mean ± SEM. Exact P values are shown.
